# Quantitative microinjection of *Candida albicans* in zebrafish embryos for infection monitoring and survival analysis

**DOI:** 10.64898/2025.12.12.693906

**Authors:** Álvaro J. Arana, Catarina Pimentel, Laura Sánchez, Oscar A. Lenis-Rojas

## Abstract

This protocol describes a versatile in vivo zebrafish embryo infection model with Candida albicans that enables precise microinjection and monitoring of defined infection readouts to compare experimental conditions (e.g., injection site and incubation temperature). The core workflow consists of quantitative microinjection of DiI-labeled C. albicans into the hindbrain ventricle of 30-48 hpf embryos, incubation at 23 deg C, and daily survival scoring to evaluate condition-dependent mortality. We calibrate the inoculum by hemocytometer counts and by plating the same injection suspension to determine colony-forming units (CFU) per pulse, allowing CFU per embryo to be reported and experiments to be compared across laboratories. Survival is analyzed with Kaplan-Meier curves and, when groups are compared, Cox models. DiI fluorescence verifies successful delivery, excludes mis-injections, and permits semi-quantitative burden comparisons between injection sites and temperatures at 24-48 hpi; beyond ∼48 hpi, signal becomes unreliable due to dye dilution and redistribution. The protocol is designed to be reproducible without transgenic reporters or confocal microscopy and includes practical guidance on randomization, blinding, sample-size planning, and reporting.

## BEFORE YOU BEGIN

Zebrafish infection models have been previously established and validated for several fungal species (Rosowski et al., 2018; Brothers et al., 2011; Hoeksma et al., 2019), providing a foundation for this protocol. The following protocol describes infection of zebrafish (*Danio rerio*) embryos with *Candida albicans* to evaluate infection dynamics and survival across experimental conditions in vivo. The procedures outlined are adapted for high reproducibility across different zebrafish laboratories and are compatible with both wild-type and non-transgenic lines, using basic fluorescence microscopy setups such as the Nikon AZ100. This makes the model especially suitable for resource-limited environments or for labs lacking access to transgenic fluorescent lines or confocal imaging (Rosowski, 2018).

This method builds upon established fungal infection models in zebrafish, including protocols for *Rhizopus arrhizus* (Wurster et al., 2021), *Aspergillus fumigatus* (Thrikawala & Rosowski, 2022), and *Candida albicans* (Brothers & Wheeler, 2012), as well as chemical approaches using fungal secondary metabolites (Hoeksma et al., 2019). Unlike systems that require epithelial damage or genetically modified lines (e.g., GAL4-UAS/NTR-based ablation strategies in Eisenhoffer et al., 2017; Atieh et al., 2021), this protocol allows infection tracking and survival assessment with minimal equipment and can be applied to a wide range of fungal isolates.

We use DiI labeling to enable tracking of *C. albicans* without genetically modified strains; because lipophilic dyes can dilute or redistribute, we include validation steps and dye-transfer controls (Nicola et al., 2009; Cseresnyes et al., 2020). Hindbrain ventricle (HBV) microinjection and multi-day monitoring follow widely used larval zebrafish protocols (Thrikawala & Rosowski, 2020; Schoen et al., 2021). All experiments complied with regulations and BSL-2 guidance. A core goal is to provide precise inoculum delivery and a reproducible survival window to compare injection site and temperature.

### OBSERVARTION NOTE (empirical differences by site and temperature)

In our hands, different combinations of injection site and incubation temperature yielded distinct infection behaviors. HBV at 23 °C typically produced a localized focus that persisted for several days, whereas yolk at 28 °C was associated with rapid dissemination and earlier mortality. These observations are detailed in Expected outcomes and can guide the choice of conditions for a given experiment.

### NOTE (fluorescence window)

In this protocol, DiI is a reliable readout from 0–6 hpi (to confirm inoculation) and remains informative at 24 and 48 hpi to discriminate differences caused by injection site (HBV vs yolk sac) and incubation temperature (23 °C vs 28 °C). After ∼48 hpi, DiI signal often fades or becomes patchy due to dye dilution and redistribution, so later images are documented but not used for quantitative comparisons.

### Recommended baseline condition

For reproducible infection monitoring across laboratories, we recommend microinjecting DiI-labeled *Candida albican* sinto the HBV of zebrafish embryos at 30–48 hours post-fertilization and maintaining infected embryos at 23 °C. This setup produces a localized infection that progresses over tens of hours without causing immediate lethality and provides a longer observation window to compare conditions.

Yolk-sac injection and incubation at 28 °C are included as optional variants. These alternative conditions drive faster systemic dissemination and earlier death. They are valuable for studying pathogenesis and virulence under high-stress conditions but are less suitable when a long observation window is required.

NOTE: All experimental procedures were conducted under BSL-2 containment and in compliance with national and European regulations on the use of animals for scientific purposes. Zebrafish husbandry, microinjection and infection procedures were approved by the Animal Care and Use Committee of the University of Santiago de Compostela and adhere to Directive 2010/63/EU and Spanish Royal Decree 53/2013.

## KEY RESOURCES TABLE

| Resource | Source | Identifier |
| --- | --- | --- |
| *Candida albicans* strain | doi: 10.3389/fmicb.2015.01382 | ASCa1 |
| RPMI-1640 | Gibco (Thermo Fisher) | 11875 (or 11875-093) |
| DiI (lipophilic tracer) | Invitrogen (Thermo Fisher) | D3911 |
| PVP40 | Sigma-Aldrich | P0930 |
| Tricaine (MS-222) | Sigma-Aldrich | A5040 |
| E3 salts (NaCl, KCl, CaCl<sub>2</sub>, MgSO<sub>4</sub>) | Sigma-Aldrich or equivalent | NaCl S7653, KCl P9541, CaCl<sub>2</sub> C1016, MgSO<sub>4</sub> M7506 |
| Low-melt agarose | Invitrogen (Thermo Fisher) | UltraPure™ LMP Agarose 16520-100 |
| YPD agar powder | Sigma-Aldrich | Y1500 |
| Petri dishes, sterile 90 mm (polystyrene, vented lids) | Thermo Scientific Nunc (or equivalent) | (specify catalog used) |
| 24-well plates, tissue-culture treated | Thermo Scientific BioLite | 130186 |
| Neubauer Improved hemocytometer | Marienfeld Superior | 0630030 (bright line) / 0630010 (dark line) |
| Borosilicate capillaries, 1.0 mm OD / 0.78 mm ID, with filament | Sutter Instrument | BF100-78-10 |
| Pneumatic microinjector | Narishige | IM-300 (or IM-31; specify) |
| Micromanipulator | Narishige | MM-3 (specify model used) |
| Mineral oil (light) | Sigma-Aldrich | M5904 |
| Stage micrometer (1 mm / 0.01 mm) | AmScope / Graticules or equivalent | e.g., MR095 or PS8 |
| Fluorescence stereomicroscope (DiI/TRITC) | Nikon (e.g., AZ100) | TRITC/DiI filter set (specify) |
| Image analysis software | NIH | Fiji (ImageJ) 2.16.0 / 1.54p |
| R (survival/coxph) | R Foundation / CRAN | R ≥ 4.2; survival package |

## MATERIALS AND EQUIPMENT

- Sterile tubes and pipette tips (1.5 mL, 15 mL, 50 mL)
- Sterile 1× PBS
- RPMI-1640 medium
- YPD agar powder (for plates)
- Petri dishes for pouring YPD plates
- Stage micrometer (1 mm / 0.01 mm) for droplet diameter measurement and volume estimation
- Embryo medium (E3) recipe (per liter): 5.0 mM NaCl, 0.17 mM KCl, 0.33 mM CaCl_2_, 0.33 mM
- 5% PVP40 in PBS (prepare fresh weekly)
- Tricaine working solution: 160 mg/L (0.016%) MS-222 in embryo medium (prepare fresh each session)
- Tricaine stock: 4 mg/mL aliquots (frozen). Dilute into the working solution as above
- DiI stock (per supplier instructions)
- Neubauer Improved hemocytometer with coverslips
- Borosilicate glass capillaries 1.0 mm OD / 0.78 mm ID with filament (for injection needles)
- Pneumatic microinjector (e.g., Narishige IM-300 or IM-31)
- Micromanipulator (e.g., Narishige MM-3; specify model used)
- Mineral oil (light) for volume calibration by droplet in oil
- Low-melt agarose 0.8% in embryo medium (E3) for mounting (use this concentration consistently)
- 24-well plates, tissue-culture treated (Thermo Scientific BioLite, Cat. 130186)
- Fluorescence stereomicroscope with TRITC/DiI filter set (e.g., Nikon AZ100)

## STEP-BY-STEP METHOD DETAILS

### 1. Preparation and DiI Labeling of *C. albicans*

**Timing:** 1–2 days

1.1. Streak the working *C. albicans* strain (ASCa1) on YPD agar plates and incubate at 30 °C, 24 h.

1.2. Inoculate a single colony into 5 mL RPMI, incubate overnight with agitation at 25 °C.

1.3. Harvest 1 mL culture (600 × g, 5 min). Resuspend in 1 mL PBS, add DiI 1% v/v, and incubate 25–30 min in the dark.

**CRITICAL STEP**. Protect DiI-labeled cells from light before and after labeling.

1.4. Count cells with a hemocytometer and adjust to 10–50 cells/nL. For each injection batch, record: (i) cells per nL (hemocytometer), (ii) viable CFU per nL (see 1.6–1.8), and (iii) initial DiI□ foci per embryo at 0–6 hpi. These three measures define the delivered inoculum.

1.5. Wash twice in PBS. Resuspend in 5% PVP40/PBS (injectable suspension).

1.6. Prepare serial dilutions (10^−4^–10^−6^) of the final injection suspension. Plate on YPD, 30 °C for 24 h to quantify CFU/mL.

1.7. To estimate CFU per pulse, eject individual 2–3 nL pulses of the final suspension directly onto

YPD (replicate injections), incubate and count colonies. **CRITICAL STEP**. Use CFU/pulse from replicate injections as the quantitative inoculum. do not infer viable dose from fluorescence intensity alone.

1.8. Compute CFU/nL = CFU/mL ÷ 10^6^. With 2–3 nL per pulse, target ∼50–100 CFU/embryo.

**RECOMMENDATION**. Gently vortex/flick the suspension before and during injections to avoid clumping/sedimentation. re-trim the needle if flow changes.

**OPTIONAL**. Assess viability with trypan blue if an additional live/dead control is required.

**NOTE (fluorescence rationale)**. DiI is robust on a standard stereomicroscope. GFP fluorescence is often too weak to visualize clearly in vivo with standard stereomicroscopes.

### 2. Microinjection into Zebrafish Embryos

**Timing:** 2–3 h

2.1. Maintain embryos (e.g., AB line) at 28 °C until 30–38 hpf.

2.2. Anesthetize in tricaine 160 mg/L (∼0.016%), pH ∼7.0, ≤5 min.

2.3. Mount in 0.8% low-melt agarose in E3, orienting the target site.

2.4. Using pulled borosilicate capillaries and a calibrated microinjector, inject 2–3 nL DiI-labeled

C. albicans into:

- HBV
- Yolk sac

2.5. Calibrate volume before starting: expel droplets into mineral oil, measure diameter with a stage micrometer, estimate V = 4/3·π·r^3^ (≈1 nL at ∼125 µm diameter), and adjust pressure/pulse for 2–3 nL.

2.6. Release embryos from agarose and transfer to fresh E3. Incubate under:

- 23 °C after HBV injection
- 28 °C after yolk injection

**OBSERVATION (typical phenotypes)**. Across experiments using matched inoculum calibration and imaging settings, HBV at 23 °C generally remained localized over multiple days, while yolk at 28 °C showed early systemic spread. Keep the site–temperature combination constant within a comparison to avoid confounding.

**CRITICAL STEP**. Injection site and temperature dictate infection spread. HBV at 23 °C yields multi-day windows for condition comparisons. Yolk at 28 °C often produces rapid, near-uniform lethality.

**RECOMMENDATION**. Perform HBV injections before 48 hpf for consistent anatomy and easier access.

### 3a. Monitoring of Fungal Proliferation

**Timing:** 1–2 h per imaging session

3a.1. 2–6 hpi imaging. Acquire low-magnification fluorescence images. Manually count discrete DiI? foci per embryo to confirm inoculum and establish a per-embryo baseline.

**CRITICAL STEP**. Use the 2–6 hpi check as an injection-quality filter. Exclude embryos with no/ectopic/weak DiI at the target site.

3a.2. 24 and 48 hpi imaging. For each condition, acquire images with identical microscope/zoom/exposure/gain/filters used at 0–6 hpi. Capture one image per embryo and record:(i)localized vs disseminated pattern, (ii) visible increase of DiI-positive area, (iii) newly involved territories. Representative patterns across conditions are shown in Figure 2A.

**Figure 1.**
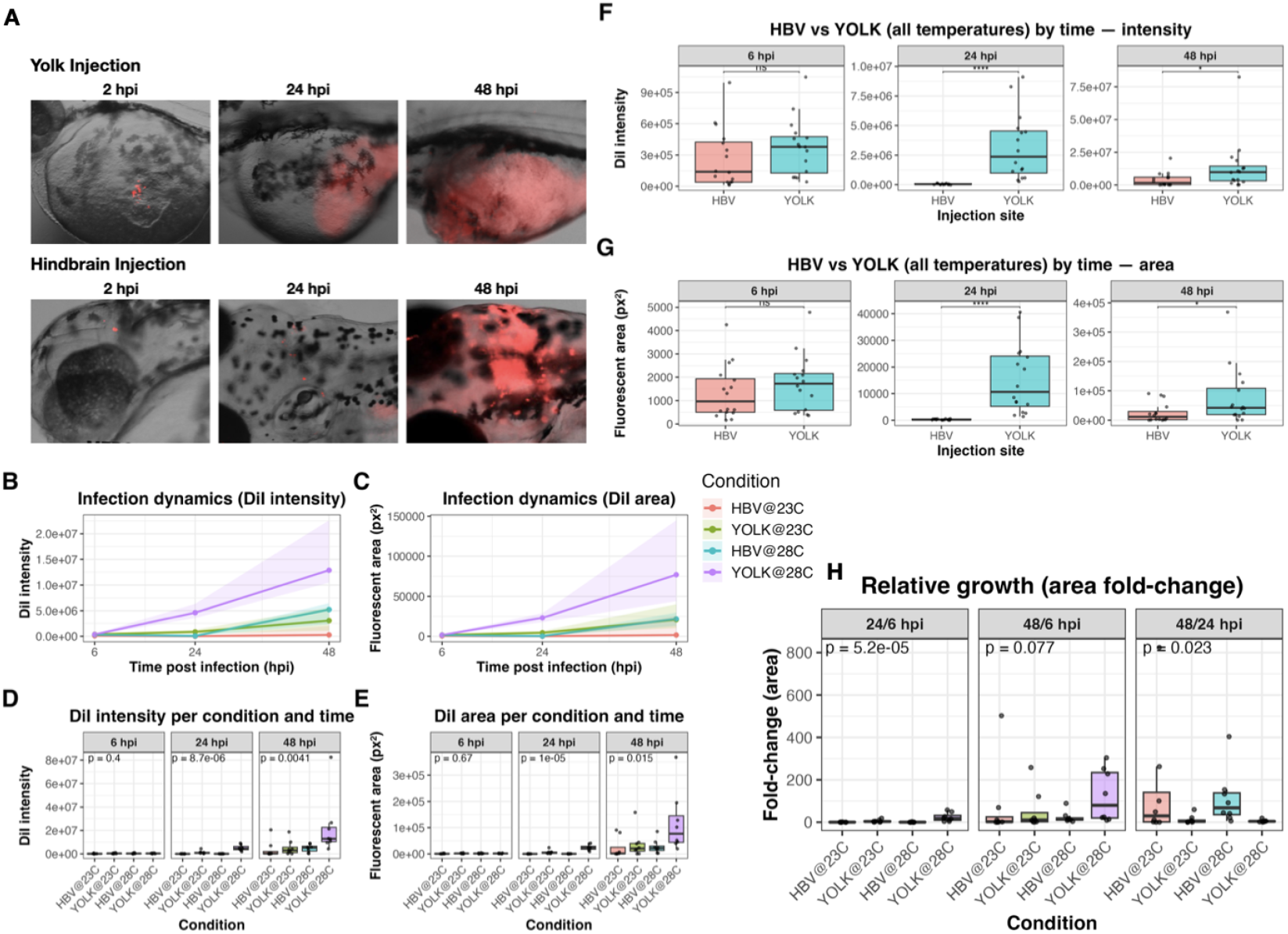
Early infection dynamics by injection site and temperature. (A) Representative bright-field/fluorescence composites at 2, 24 and 48 hpi into the yolk (top, YOLK) or the hindbrain ventricle (bottom, HBV) at 28 °C. DiI signal (red) marks the inoculum. Time-course of (B) total DiI intensity or (C) fluorescent area (pixels) at 6, 24 and 48 hpi for the four conditions (HBV at 23 °C, Yolk at 28 °C, HBV at 28 °C, Yolk at 28 °C). Lines show the mean; shaded bands show variability. Boxplots of (D) DiI intensity and (E) fluorescent area by condition at each time; the panel header reports the global *p*-value. Kruskal–Wallis across all groups at each time; when applicable, Dunn’s post-hoc (Benjamini–Hochberg adjusted. (F) HBV vs YOLK (temperatures pooled) at 6, 24 and 48 hpi for DiI intensity. tTwo-sided Wilcoxon rank-sum per time; *p*-values shown on each facet. (G) HBV vs YOLK (temperatures pooled) for fluorescent area at 6, 24 and 48 hpi. Wilcoxon rank-sum per time; *p*-values shown. (H) Relative growth of fluorescent area (fold-change 24/6, 48/6 and 48/24 hpi) across conditions; global *p*-value in each facet. Kruskal–Wallis + Dunn’s post-hoc when needed. Each dot is one larva; boxes show median and IQR. Significance is reported as numeric *p*-values on the plot; “ns” denotes *p* ≥ 0.05. Abbreviations: HBV, hindbrain ventricle injection; YOLK, yolk-sac injection; hpi, hours post infection; DiI, lipophilic tracer.

**Figure 2.**
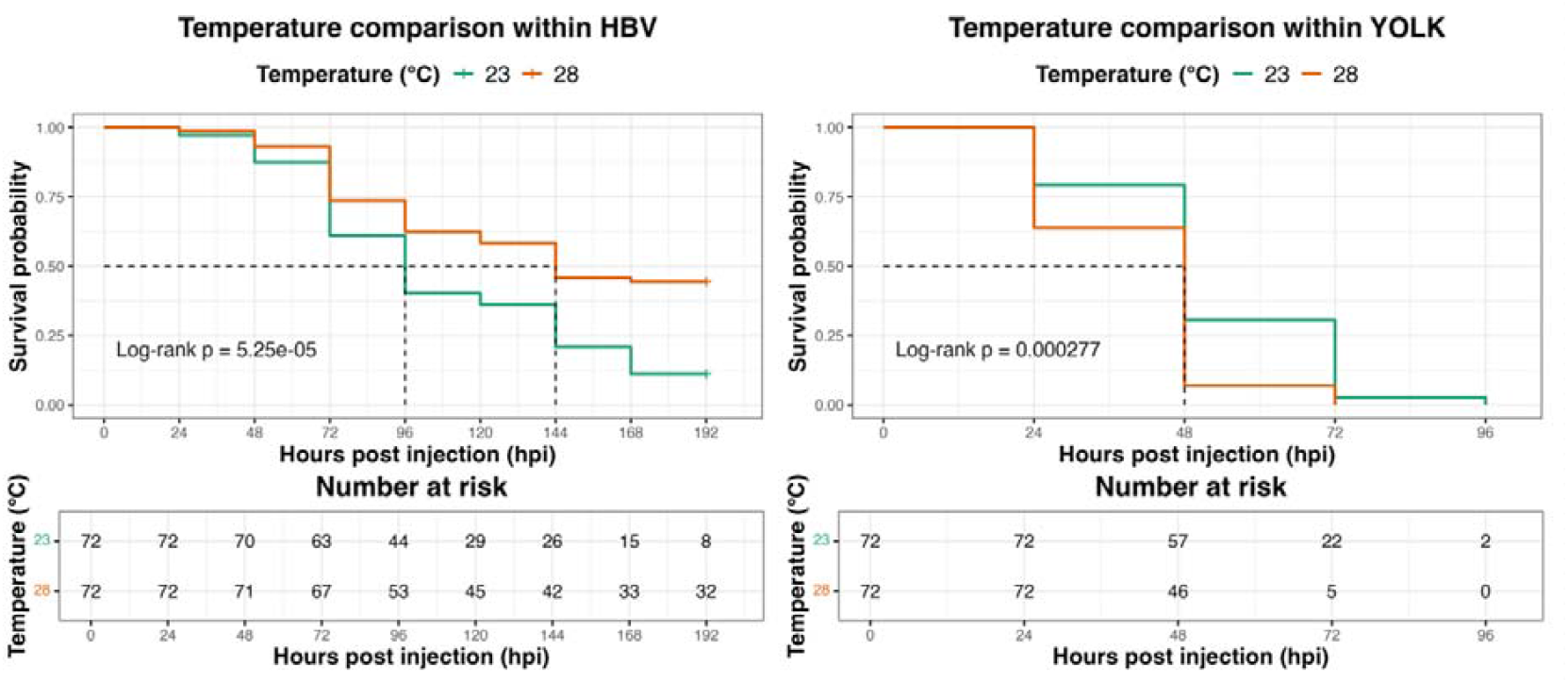
Temperature-dependent survival after Candida albicans infection in zebrafish larvae. Kaplan–Meier curves comparing incubation at 23 °C and 28 °C following microinjection into (A) the HBV, monitored up to 8 dpi (192 hpi), or (B) the yolk, monitored up to 4 dpi (96 hpi). Dashed lines indicate 50% survival; “+” marks censored individuals, and tables show the number at risk. Survival distributions were compared using the log-rank (Mantel–Cox) test: HBV, p = 5.25 × 10□□; yolk, p = 2.77 × 10□□. n = 72 larvae per temperature group (per injection site).

**NOTE**. Images at 24/48 hpi can be semi-quantitatively compared within an experiment because acquisition settings are identical.

**LIMITATION**. Do not extend these comparisons beyond 48 hpi. DiI becomes non-uniform and no longer reflects total burden.

3a.3. Fiji-based quantification (6–48 hpi).

1. Open the image in Fiji. convert to 8-bit if needed.
2. Apply a single predefined threshold (e.g., Default or Moments) selected on pilot images from the same setup.
3. Draw a fixed ROI for the infected compartment (HBV or yolk) and reuse for all embryos in that experiment.
4. Measure area (px^2^ or µm^2^) and integrated density of DiI-positive signal.
5. Export to CSV and label: experiment ID, embryo ID, injection site, temperature, time (6, 24, 48 hpi).

Example quantification outputs are shown in Figure 2B–H.

**EXPECTED OUTCOME**. 24/48 hpi values discriminate faster/disseminated (Yolk at 28 °C) from slower/localized (HBV at 23 °C) infections under matched imaging.

**DO NOT**. (i) Pool images from different days/scopes. (ii) run multi-timepoint tests across batches.(iii) treat >48 hpi DiI as quantitative (see Figure 2).

3a.4. Representative images. Document examples of localized (HBV at 23 °C) and disseminated (Yolk at 28 °C) infections to illustrate pattern-by-condition.

**INTERPRETATION NOTE**. DiI intensity per cell decreases as yeast divide. Hyphal growth increases apparent area. Treat DiI as a semi-quantitative descriptor of spread/morphology under matched settings, not as absolute burden.

### 3b. Absolute fungal burden per larva (CFU/larva)

**Timing:** 1–2 h per batch

3b.1. Euthanize ≥10 larvae per condition at defined times (e.g., 6, 24, 48 hpi) with ice-cold tricaine overdose.

3b.2. Rinse briefly in sterile PBS. Transfer each larva to 200 µL PBS in a microtube with a sterile pestle.

3b.3. Homogenize thoroughly. Prepare serial dilutions (10^−1^–10^−3^). Plate on YPD (add antibiotics if needed).

3b.4. Incubate 24–48 h at 30 °C and count colonies for CFU/larva.

3b.5. Report median [IQR] per condition/time. normalize to initial inoculum (CFU per 2–3 nL pulse).

**NOTE**. In routine use, define inoculum a priori as CFU/nL from replicate plated pulses (Section 1), and record early DiI^+^ foci (Section 3a) as a fast quality control.

### 4. Survival Scoring and Data Analysis

**Timing:** daily, up to planned endpoint (e.g., 5–8 dpi)

4.1. Daily scoring. Assess viability without anesthetics. verify heartbeat/circulation under a stereomicroscope.

**CRITICAL STEP**. Do not use tricaine during survival checks. it can transiently suppress visible heartbeat and cause false death calls.

4.2. Set pre-specification. Before starting, define: primary endpoint (e.g., mortality by 5 or 8 dpi), randomization, blinding (where feasible), and what counts as biological replicate (clutch/experiment) and technical unit (well).

4.3. Analysis. Plot Kaplan–Meier curves for each condition. compare with log-rank tests. Fit Cox models including condition (and design covariates if relevant) with a frailty term for well or batch. report hazard ratios with 95% CIs and adjust p-values for multiple comparisons. 4.3Example Kaplan–Meier curves for the main comparisons are shown in Figure 3.

4.4. Inclusion flow. Provide a simple diagram of embryo inclusion/exclusion (e.g., excluded at 2 hpi for failed injection/absent DiI).

**OPTIONAL (within Step 4)**. If compounds are included, analyze survival in the same Kaplan– Meier and Cox framework with matched vehicle and toxicity controls (NOINF+VEH, NOINF+COMPOUND).

**REPORTING, DATA & CODE AVAILABILITY**. Provide raw fluorescence images (original resolution), CFU count tables, injector calibration data (volume per pulse, CFU per nL), and the ImageJ/Fiji macro used for single-threshold, single-ROI measurements at 2–48 hpi, together with the R code for Kaplan–Meier and Cox models. Indicate that Fiji measurements are valid only for comparisons within the same experiment and up to 48 hpi.

## EXPECTED OUTCOMES

Under the recommended baseline condition (HBV injection at 30–48 hpf followed by incubation at 23 °C), infected embryos typically develop a progressive but initially localized *Candida albicans* infection that persists for several days without immediate, nonspecific lethality. This produces a reproducible observation window in which survival can be compared across conditions such as injection site and incubation temperature using Kaplan–Meier curves and Cox models with a shared frailty term for well or batch. Effect sizes can be reported as hazard ratios with 95% confidence intervals for the relevant group contrasts.

At later time points (24 and 48 hpi), DiI imaging allows semi-quantitative discrimination between slower, localized infections (HBV at 23 °C) and faster, disseminated infections (yolk at 28 °C), provided that images are acquired with identical settings on the same microscope. After 48 hpi, DiI often becomes faint or patchy and should not be used for quantitative comparisons, although it remains useful to illustrate overall infection patterns.

Absolute fungal burden can be measured directly as CFU/larva by homogenizing euthanized larvae at defined time points (for example 6, 24, 48 hpi), plating serial dilutions on YPD, and counting colonies. In routine practice, the inoculum is defined a priori as CFU per nL by plating replicate injection pulses of the exact DiI-labeled suspension, and the initial number of DiI□ foci per embryo at 0–6 hpi is recorded as a quality control. These linked measurements (CFU/nL, CFU per 2–3 nL pulse, early DiI□ foci) support comparisons across embryos, wells, and laboratories.

For reference, yolk-sac injection at 28 °C yields a high-virulence phenotype characterized by rapid systemic dissemination and earlier death. This condition is valuable for studying dissemination kinetics and virulence under high stress, but because mortality can become near-uniform early, it is generally not optimal when an extended observation window is required.

Representative output for publication should include: (i) Kaplan–Meier survival curves with hazard ratios and confidence intervals for the main group comparisons; (ii) example fluorescence images at 6–48 hpi illustrating localized versus disseminated infections; (iii) CFU data or inoculum calibration tables; and (iv) full analysis scripts and raw image datasets to document reproducibility (see Figure 3).

## LIMITATIONS

This protocol depends on precise and consistent microinjection technique. Delivering ∼2–3 nL into anatomically small sites such as the HBV requires practice and volume calibration. Small deviations in injection depth, angle or volume can lead to substantial differences in effective inoculum between embryos, which in turn affects downstream survival curves and hazard-ratio estimates. Calibrating injection volume in mineral oil and excluding embryos that lack clear DiI□ inoculum at 2 hpi are essential steps to control this source of variability.

The optical readout is constrained by the behavior of DiI. DiI provides a strong initial signal that allows to (i) confirm delivery of the inoculum and (ii) follow qualitative spread and morphology over the first 24–48 hpi. However, DiI is a lipophilic membrane dye that dilutes as *Candida* divides and redistributes along elongating hyphae. As a result, fluorescence intensity per structure is not stable over time, and apparent “burden” can increase either because the fungus proliferates or because dye spreads along hyphal branches. Photobleaching and dye transfer into host material or debris can further complicate interpretation (Nicola et al., 2009; Cseresnyes et al., 2020). For that reason, DiI signal at later time points should be treated as semi-quantitative and always compared under strictly matched imaging settings and time points, rather than as an absolute fungal load per larva. When precise burden is required, direct CFU/larva measurements from homogenized embryos are still necessary.

Finally, this assay is optimized for gross infection dynamics and survival analysis, not cellular-resolution host–pathogen biology. It is intentionally designed to run in non-transgenic embryos on a standard fluorescence stereomicroscope rather than requiring confocal imaging or fluorescent reporter fish. This makes it portable and scalable, but it also means that detailed questions about fungal uptake by specific immune subtypes, intracellular residency, or subcellular host damage may require complementary approaches using transgenic reporters, higher numerical aperture objectives, or confocal microscopy.

## TROUBLESHOOTING

### Problem 1

No DiI signal observed after microinjection.

This most commonly reflects incomplete/uneven DiI labeling, loss of signal due to unnecessary light exposure before imaging, sedimentation of yeast in the needle (very few labeled cells delivered), or failed delivery (missed compartment / leakage).

### Potential solution

Use freshly prepared DiI-labeled *Candida* and keep the suspension protected from light before and after staining. Mix the inoculum immediately before loading the needle and periodically during injections to prevent settling. Minimize light exposure during screening and acquire images promptly. If the issue persists, verify that labeled cells are clearly fluorescent before starting the session, and confirm correct delivery by screening embryos early (e.g., ∼2 hpi) to ensure a clear DiI□ bolus at the intended site.

### Problem 2

High mortality in all injected and control groups.

Generalized death in both infected and control larvae usually results from excessive injection volume, traumatic needle insertion, contaminated carrier/buffer solutions, or anesthesia issues (incorrect tricaine dilution and/or pH).

### Potential solution

Calibrate injection volume in mineral oil and keep pulse volume within the protocol range (avoid over-injection). Standardize needle angle and depth to minimize mechanical trauma. Prepare carrier/buffer and tricaine fresh, use sterile technique, and discard any solution that appears cloudy. Confirm tricaine working concentration and pH as specified in the protocol to avoid anesthetic overdose or stress.

### Problem 3

Inconsistent infection outcomes across embryos.

Some embryos show strong fungal expansion while others appear essentially uninfected. This typically reflects off-target injections (wrong compartment), variable delivered volume, partial needle clogging, or embryos that did not receive viable inoculum.

### Potential solution

At early QC (e.g., ∼2 hpi), screen embryos by fluorescence and keep only those with a clear DiI□ inoculum in the intended compartment. Exclude embryos with absent signal or signal in the wrong location before randomization/treatment allocation. Re-trim or replace the needle when flow becomes inconsistent, and re-check volume calibration after any needle adjustment.

### Problem 4

Weak or fading DiI signal at later time points (48 hpi and beyond).

Apparent signal loss can reflect photobleaching across repeated imaging, dilution/redistribution of the lipophilic dye as the fungus grows, and changes in fluorescence appearance during hyphal development. This can make late images look “weaker” even when fungus is still present.

### Potential solution

Limit light exposure during imaging and keep larvae in the dark between sessions. Use constant exposure time, gain, and magnification across time points. Treat late DiI images as qualitative pattern documentation, not an absolute burden readout. If absolute fungal burden is needed at later stages, switch to CFU-per-larva measurements from homogenized embryos rather than relying on DiI intensity alone.

### Problem 5

Apparent DiI signal in host-derived structures (dye redistribution).

Because DiI is membrane-associated, fluorescence can appear in debris or host-derived material during clearance, which can be misinterpreted as “live fungus,” especially at later time points.

### Potential solution

Do not assume that all late DiI signal represents viable *Candida*. Interpret fluorescence together with morphology and with the early QC images confirming correct delivery. When interpretation is uncertain (especially ≥48 hpi), confirm outcomes **using** CFU-per-larva rather than relying on DiI signal alone.

### Problem 6

GFP fluorescence is not visible in vivo.

With GFP-reporter *Candida* strains, GFP can be difficult to detect in live embryos on standard fluorescence stereomicroscopy due to embryo autofluorescence, light scattering, weak in vivo signal, incorrect filter set, or photobleaching.

### Potential solution

Confirm you are using a GFP/FITC filter set and appropriate illumination, and increase exposure/gain within non-saturating limits. If GFP remains undetectable, proceed with DiI-based tracking (the standard readout in this protocol) and confirm strain identity ex vivo (e.g., on plates) or in fixed samples using a more sensitive imaging setup if available.

## ACKNOWLEDGMENTS

We are grateful to Prof. Anna Krasowska for providing the *Candida albicans* ASCa1 strain.This work was supported by FCT - Fundação para a Ciência e a Tecnologia, I. P., through MOSTMICRO-ITQB R&D Unit (UIDB/04612/2020, UIDP/04612/2020), LS4FUTURE Associated Laboratory (LA/P/0087/2020) and the IC&DT FCT project LISBOA2030-FEDER-00693800 “*Study and optimization of a novel, first-of-its-kind, marine-derived antifungal drug” (to CP)*.

## Graphical Abstract

Workflow for zebrafish embryo *Candida albicans* infection. Step 1 – Staining. DiI labeling of *C. albicans* to track the inoculum. Step 2 – Microinjection. Pneumatic injection of labeled yeast (2–3 nL) into the yolk sac or the HBV of zebrafish embryos. Step 3 – Infection monitoring. Fluorescence microscopy at predefined time points (e.g., 6 and 48 hpi) to visualize and quantify fungal signal. Step 4 – Survival analysis. Daily survival scoring. Analyze data with Kaplan–Meier curves (log-rank/Mantel–Cox where applicable). Intermediate step (optional) – Absolute CFU.(a) Plate serial dilutions of the injected suspension; (b–c) dissociate single embryos at selected time points and plate to enumerate CFU. Grey dots mark sampling/analysis points; red-circled letters (a–c) correspond to CFU panels. Abbreviations: HBV, hindbrain ventricle; hpi, hours post-infection; CFU, colony-forming units.

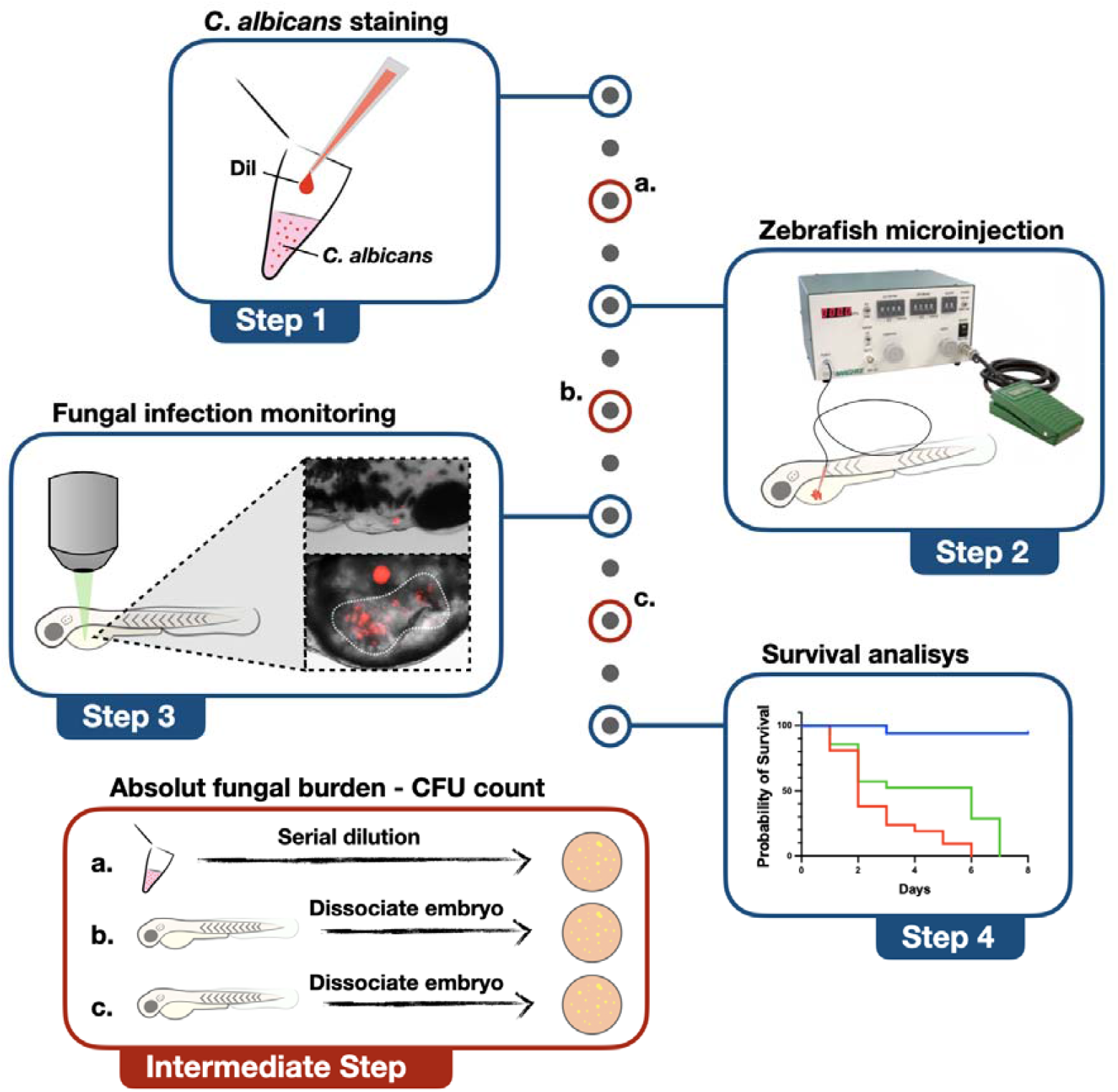

